# *Leishmania tarentolae* and *Leishmania infantum* in geckos from Mallorca Island, Spain

**DOI:** 10.1101/2025.02.06.636881

**Authors:** Joan Martí-Carreras, Johan Espunyes, Laura Carrera-Faja, Carlotta Pasetto, Maria Magdalena Alcover Amengual, Sarah Chavez-Fisa, Marina Carrasco, Xavier Roura, Olga Francino, Lluís Ferrer

## Abstract

*Leishmania tarentolae* and *Leishmania infantum* are two sympatric parasites of significant ecological and epidemiological interest in the Mediterranean basin. This study investigated the prevalence of *L. tarentolae* and *L. infantum* in two gecko species (*Tarentola mauritanica* and *Hemidactylus turcicus*) present on Mallorca Island, Spain, using duplex quantitative PCR. A total of 59 geckos were sampled across the island, including 53 *T. mauritanica* and six *H. turcicus*. Tissue and blood samples were screened for both parasites, and generalized linear models were used to assess organ-specific infection patterns. The results revealed the prevalence of *Leishmania* infection in adult *T. mauritanica*, with 10/49 (20.41%) testing positive for *L. tarentolae* and with 1/49 (2.04%) for *L. infantum*. Coinfection with both parasites was detected in 3/49 geckos (6.12%). No positives were identified in *H. turcicus*, probably due to small sample size. Regarding positivity by tissues, coleomic organs were more likely to be positive for *L. tarentolae* in adult *T. mauritanica* than blood, with a slighter positivity in the liver, spleen and lung. Interestingly, *L. infantum* was detected more frequently coinfecting with *L. tarentolae* than as a single parasite. These prevalence rates and the existence of coinfections highlight potential ecological interactions between the two parasites and reptiles. This study provides valuable data on the potential role of geckos in endemic areas like Mallorca.

**Author Summary:** Leishmaniosis is a zoonotic disease caused by parasites that can infect both animals and humans. In Europe, *Leishmania infantum* is the main species responsible for the disease, with dogs acting as its primary host. However, recent research suggests that other animals, including reptiles, might also play a role in their spread. In this study, centered in wild geckos from Mallorca Island (Spain), we investigated the prevalence of *L. tarentolae* and *L. infantum* in two gecko species (*Tarentola mauritanica* and *Hemidactylus turcicus*). Using quantitative PCR, it was found that 14/49 (28.57%) of adult *Tarentola mauritanica* were positive for *Leishmania* parasites, 13/49 (26.53%) for *L. tarentolae* and 74/49 (8.16%) for *L. infantum*. Additionally, coinfection of both parasites was detected in three geckos (3/49, 6.12%). The second gecko species studied, *Hemidactylus turcicus*, was not positive for *Leishmania*, possible due to a small sample size. These results reinforce previous studies suggesting that geckos may carry both parasites. Since *L. tarentolae* has also been detected in dogs, and both *Leishmania* species are sympatric, it raises questions about the role of reptiles in leishmaniosis cycle and transmission, as well as the reliability of current diagnostic tests in coinfections of closely related parasite species.

## Introduction

In Europe, zoonotic visceral leishmaniosis is caused by *Leishmania infantum*, dogs being their primary reservoir. This disease has a wide distribution throughout the Mediterranean basin (1,2). *L. infantum* and *Leishmania donovani* are the dominant pathogenic species of *Leishmania* present in western and central Europe (3). In recent decades, different studies have shown that the epidemiology of leishmaniosis in Europe is much more complex than previously thought, broadening the range of reservoir hosts species (4) or reporting an increase in drug resistance (5–7). *Leishmania infantum* infections have been detected in other species of vertebrates, such as lagomorphs (i.e., rabbits and hares), which play an important role as sylvatic reservoirs in endemic areas of Europe (8–11). More recently, the infection of different Mediterranean reptile species with *L. infantum* has also been demonstrated, although its epidemiological role remains to be elucidated in this case (12).

Among Mediterranean reptiles, the most frequently reported *Leishmania* species is *Leishmania tarentolae*. This species, subgenus *Sauroleishmania*, is limited to the Old World and it is found in association with geckos in southern Europe and North Africa (13,14), and possibly also other areas with endemic Old World gecko populations, such as the Middle East (15). *Leishmania tarentolae* is transmitted by reptile-biting sandflies of the genus *Sergentomya* and is considered sympatric with *L. infantum*. It is regarded as non-pathogenic to mammals, including dogs (3,16,17), yet infection and parasite persistence can occur (18). Studies in Italy have detected infection rates of *L. tarentolae* up to 7% in geckos and 16.5% in lacertids (12,14). In Spain, where *Sergentomya minuta* is widely distributed (19) and where canine leishmaniosis shows high prevalence rates, especially in the Balearic Islands (20–22), it is under research which *Leishmania* species infect reptiles and which role they may play on leishmaniosis epidemiology and *Leishmania* transmission cycles (23).

Only two species of saurids inhabit the island of Mallorca, both of which are geckos: *Tarentola mauritanica* and *Hemidactylus turcicus* (24). There are no native representatives of the *Lacertidae* family, except for a few representatives of *Podarcis pituytensis,* recently introduced from the neighboring island of Eivissa (24). The two gecko species are very abundant throughout the island, with reported densities of over 1200 individuals/km^2^, due to the presence of a very suitable biotope and the absence of competitors (24). Both species are anthropophilic, often living in houses and gardens. It is thought that these geckos have been introduced to Mallorca from North Africa (*T. mauritanica*) and the eastern Mediterranean (*H. turcicus*) more than 2,000 years ago (24).

Considering the biology of these two gecko species and the high prevalence of canine leishmaniosis in Mallorca (20,22), the study hypothesis was that the gecko populations in Mallorca could be parasitized by both *L. tarentolae* and *L. infantum*. Therefore, the aim of this study was to assess the prevalence of *L. tarentolae* and *L. infantum* in gecko species at different locations of Mallorca Island using nucleic acid detection (qPCR), instead of serological testing, as current *L. infantum* serology may cross-react with *L. tarentolae* (16).

## Material and methods

### Field trapping and dissection

*Tarentola mauritanica* (common wall gecko, *dragó comú*) and *Hemidactylus turcicus* (Mediterranean house gecko, *dragonet* or *dragó rosat*), as illustrated in **Figure 1a**, were captured over five days in May 2024 in the northern and central regions of Mallorca Island (Balearic Islands, Spain), as seen in **Figure 1b**. Geckos were captured by hand and identified to species level using reference keys (25). They were then humanely euthanized in situ using intramuscular anesthesia consisting of ketamine (20 mg/kg) and medetomidine (0.2 mg/kg), followed by pitching, in accordance with the guidelines for the euthanasia of animals published by the American Veterinary Medical Association (26). The whole blood was immediately obtained via cardiac puncture and stored in a portable refrigerator before being frozen at −20ºC after each capture session. The geckos were dissected, and the spleen, heart, lungs, liver, and kidneys were individually collected and frozen at −20ºC for DNA extraction. For juvenile geckos with smaller body sizes, coelomic organs were not separated but frozen whole. Complete sample information is provided in **Supplementary Table 1**. The capture, euthanasia, and sampling in the present study was conducted following local law compliance on ethical and legal requirements for sampling wild animals (see **Ethical statements**).

**Figure 1.**
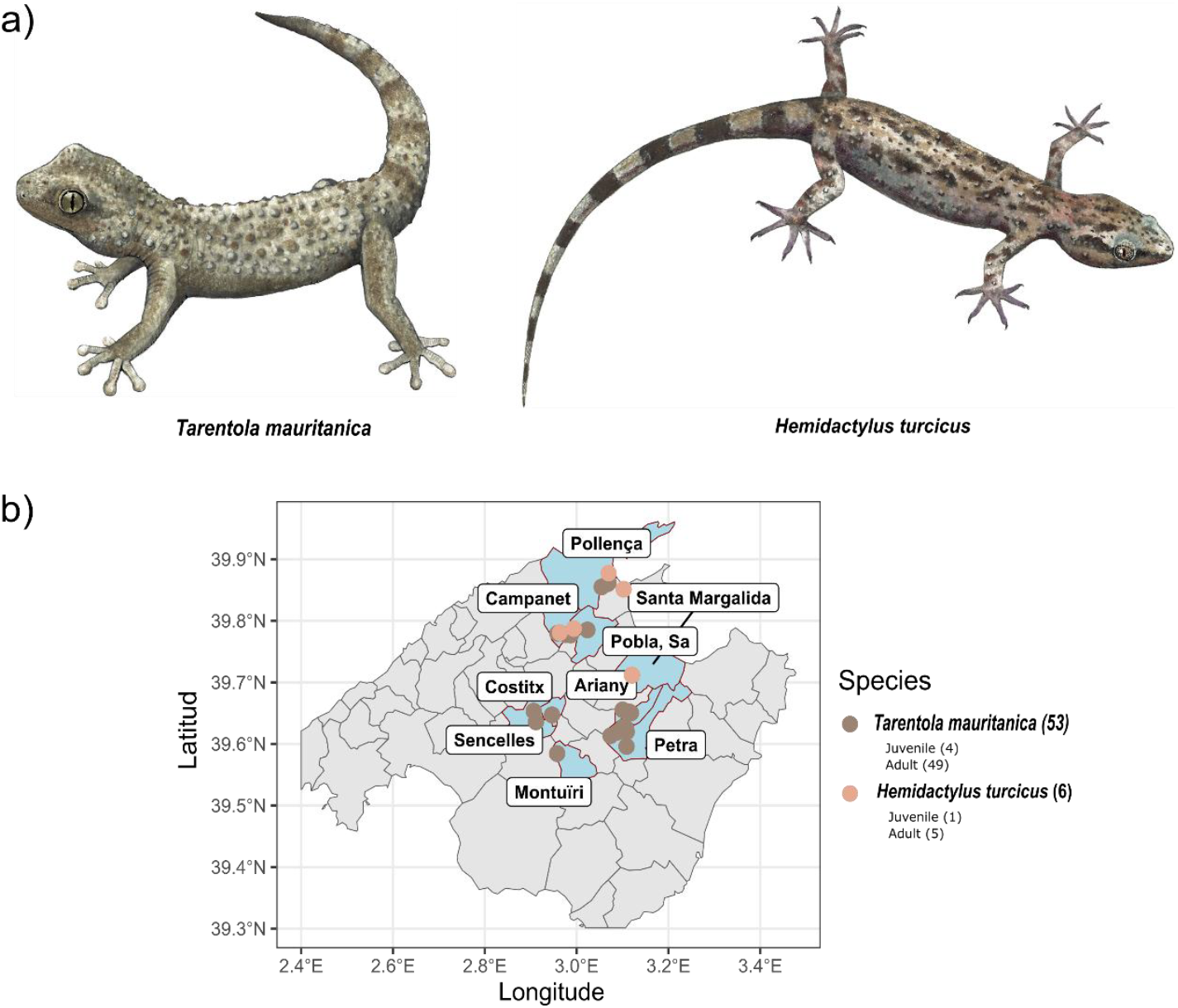
Sampling distribution of *T. mauritanica* and *H. turcicus* in Mallorca Island in May 2024. Panel a) Colored illustration of adult *T. mauritanica* and *H. turcicus* by Bruna Roqué. Panel b) Sampling distribution of *T. mauritanica* (grey) and *H. turcicus* (pink) by place of sampling (municipalities of Mallorca) and life stage (juvenile or adult).

### DNA extraction and qPCR screening

Tissue and blood samples were extracted on 96-well plates with an extraction robot Chemagic 360 (Perkin Elmer, Waltham, Massachusetts) using two different extraction protocols: Chemagic DNA Tissue10 360 prefilling H96 VD150506 for solid tissue samples and Chemagic DNA Blood 250 prefilling 360 VD180628 for blood samples. Elution was performed in 80 µl. Samples were diluted 1/10 before proceeding to qPCR screening. The TaqMan Fast Advanced Master Mix (Themo Fisher Scientific, Waltham, Massachusetts, USA) was used for screening the presence of *L. tarentolae* and *L. infantum* with a discriminative duplex qPCR system previously described targeting the ITS1 region: Fw 5’-GCAGTAAAAAAAAGGCCG-3’, *L. tarentolae* probe 6-FAM-5’-CACGCCGCGTATACAAAAACAC-3’-f1-quencher-MGB and *L. infantum* probe VIC-5’-TAACGCACCGGCCTATACAAAAGCA-3’-f1-quencher-MGB (14) with a different reverse primer Rv L5.8S-5’-TGATACCACTTATCGCACTT-3’ (27). Testing was conducted in duplicate with extraction controls for both blood and tissue samples. Positive controls for *L. tarentolae* and *L. infantum* were used. Non-template controls (NTC) were added to each qPCR plate. The analysis threshold for qPCR was set at 0.052 for *L. tarentolae* and 0.039 for *L. infantum*. A sample was considered positive if the Ct value was ≤ 38.5 for both replicates after revision of the amplification curves. Kinetoplast qPCR: LEISH1 5’-AACTTTTCTGGTCCTCCGGGTAG-3’, probe FAM-5’-AAAAATGGGT GCAGAAAT-3’ f1-quencher-MGB and LEISH2 5’-ACCCCCAGTTTCCCGCC-3’ was used for confirmation of *L. infantum* (28). Coinfection was considered when a gecko had the presence of both parasites in the same tissue sample or in different tissue samples from the same gecko. No significant amplification was identified in extraction or NTC controls.

### Data analysis

Primers and probes were evaluated using blastn (29) against the NCBI Nucleotide database (June 2024) using the recommended parameters for primer and probe design: expected threshold of 1000, word size of 7 and unmasking low complexity regions (30).

Eds qPCR files were processed with app.thermofisher (https://apps.thermofisher.com/apps/spa/), where amplification curves were manually revised. Data analysis was conducted with R v4.4.1 (31). Sample distribution was located with packages mapSpain v0.9.2, tidyverse v2.0.0 (32). Generalized linear models (binomial logit) were employed to identify which organs were statistically more prone to be positive for *L. tarentolae* and *L. infantum* with packages for logistif v1.26.0 (33) and emmeans v1.10.5 (34). Analysis was only conducted for adult *T. mauritanica* (n = 49) as juvenile *T. mauritanica* or *H. turcicus* did not have enough sample size.

## Results

A total of 59 geckos were collected, comprising five juveniles and 55 adults. Of these 59 geckos, 53 individuals were identified as *T. mauritanica* and six as *H. turcicus*, as shown in **Figure 1b**. Duplex qPCR was performed for the simultaneous detection of *L. tarentolae* and *L. infantum*. Counting both species, 14/59 (23.73%) geckos were positive for *Leishmania* spp. with 13/59 (22.03%) for *L. tarentolae* and 4/59 (6.77%) for *L. infantum*. Three of the four positives for *L. infantum* were also positive for *L. tarentolae*, thus counting as coinfection cases (**Figure 2**).

**Figure 2.**
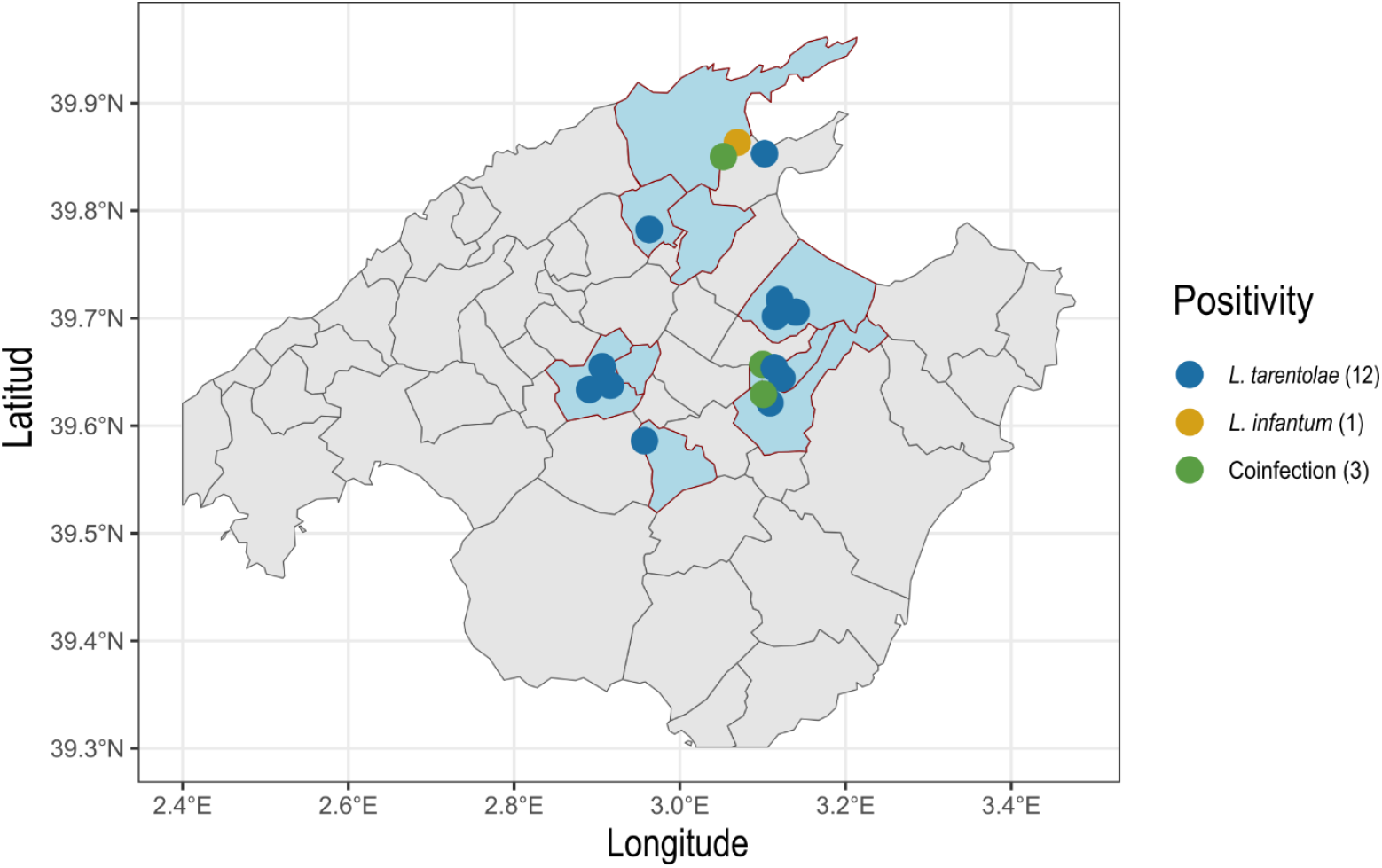
Distribution of *L. tarentolae* and *L. infantum* positive geckos in Mallorca Island in May 2024. Gecko positivity by RT-qPCR against *L. tarentolae* (blue), *L. infantum* (yellow) and coinfection with both parasites (green).

For adult *Tarentola mauritanica* (n = 49), the prevalence of *L. tarentolae* was 13/49 (26.53%) and 4/49 (8.16%) for *L. infantum*, with a coinfection rate of 3/49 (6.12%). In detail, *L. tarentolae* tested positive in 13 adults in liver (6), spleen (6), lung (6), kidney (4), heart (4) and blood (0). Regarding *L. infantum*, four adults tested positive in blood (2), spleen (2), liver (1), lung (1) and heart (1). The three coinfections detected were detected in spleen (2), liver (1), lung (1), heart (1) and blood (1), as summarized in **Table 1**. No juvenile specimens were positive for *Leishmania*. Generalized linear models were used to identify which organs were statistically more prone to be positive for *L. tarentolae* and *L. infantum* (not accounting for coinfection cases, **Table 1**) and provide guidance for their potential use for parasite detection. In adult *T. mauritanica* (n = 49), *L. tarentolae* is more likely to be identified in coelomic organs (heart, liver, spleen, lung and kidney), when compared to blood (glm, p-value = 0.0266), with a slightly higher positivity in liver, spleen and lung. Regarding *L. infantum*, no organ was significantly more likely to be positive due to its smaller sample size.

**Table 1.**
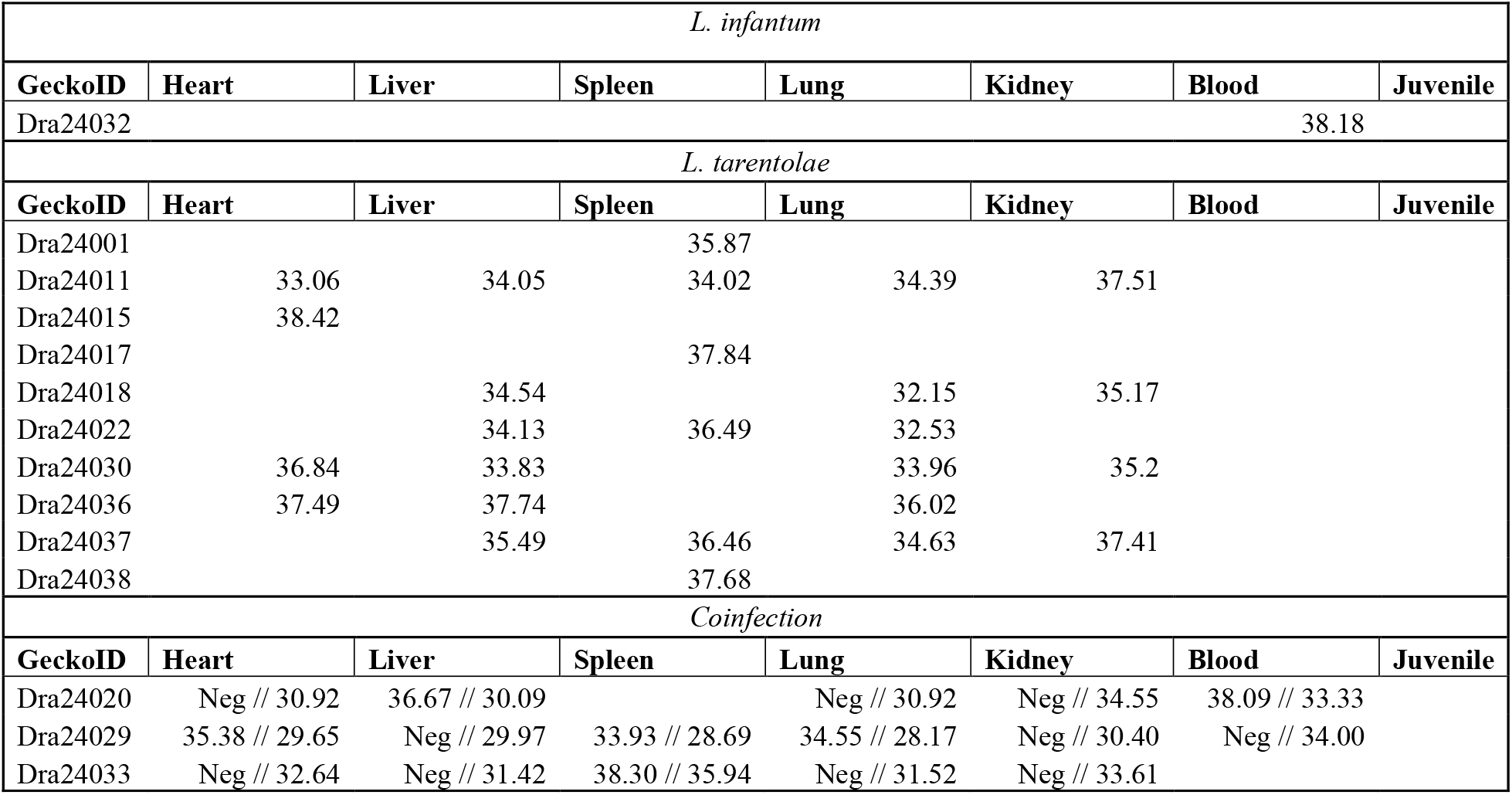
Prevalence of *L. tarentolae* and *L. infantum* by qPCR in all coelomic organs and blood in *T. mauritanica*. Detection is expressed as Ct value for positives, and Neg. for negative samples for coinfections. For each positive organ and gecko, the average Ct value is provided for *L. tarentolae* and *L. infantum*. For coinfections the average Ct is first given for *L. infantum* and after double bar to *L. tarentolae*.

For *L. tarentolae*, the average Ct of detection was 34.02 with a minimum Ct of 28.69 and maximum Ct of 38.42, meanwhile for *L. infantum*, the average Ct value of detection was 36.44 with a minimum Ct of 34.55 and a maximum Ct of 38.33. Primer and probe in silico analysis showed that TaqMan-MGB probes were highly specific against their targets. The TaqMan-MGB Probe Lt had 100% identity and 100% coverage against *L. tarentolae* (only taxon that matched). The TaqMan-MGB Probe Li had a 100% identity and 100% coverage against *L. donovani-infantum* complex. The complete specificity of TaqMan-MGB probes ensures the appropriate signal for each species. No cross hits were identified between probes, although primers align against several *Leishmania* species (*L. tarentolae, L. infantum, L. donovani, L. amazonensis, L.major, L. tropica* and *L. mexicana*) with a 100% identity and 100% sequence coverage. Additionally, positive samples for *L. infantum* Li TaqMan-MGB probe were also tested with a different qPCR assay targeting the kinetoplast for confirmation (24).

## Discussion

The Balearic Islands, including Mallorca, are endemic for *L. infantum*, the primary causative agent of leishmaniosis in humans and companion animals (20–22). In humans, the average annual incidence of clinical leishmaniosis ranges from 0.7 to 3.5 cases per 100,000 inhabitants (22). However, prevalence studies reveal that up to 5.9% of human blood donors harbor *L. infantum* DNA, indicating that most infections are asymptomatic (21). Similarly, *L. infantum* prevalence is high in dogs, with positivity ranging from 56% to 67% depending on dog population (free-living vs. kennel) or testing method (serology vs. PCR) (20,35). In cats, while data specific to the Balearic Islands is limited, the seroprevalence of *L. infantum* across Spain is approximately 15%, comparable to the Mediterranean average of 17.3% (36). It is worth noting that serological testing of *L. infantum* may be overreporting the incidence of infection, as current serological tests may not discriminate between *L. infantum* (16) and *L. tarentolae*, believed to be non-pathogenic but able to infect and persist in mammals (3,16–18).

Regarding *L. tarentolae*, its prevalence in Spain is less well-documented, as it is primarily associated with reptiles. *Leishmania tarentolae* or *L. tarentolae*-like parasites have been reported in *Sergentomyia minuta* in continental Spain (37,38). This study represents one of the most comprehensive assessments of *L. tarentolae* and *L. infantum* prevalence in wild gecko populations on Mallorca Island, investigating whether these geckos can serve as hosts for both *Leishmania* species. *T. mauritanica* exhibited notable prevalences of *L. tarentolae,* with some coinfection cases with *L. infantum* observed in this species. Regarding *H. turcicus*, no positive samples were detected, possibly due to a small sample size. In similar studies performed in the south of Italy, Latrofa *et al.*, 2021, reported no cases of *L. tarentolae* or *L. infantum* in *T. mauritanica*, albeit with only three geckos (14). Two studies from Mendoza-Roldán *et al.*, reported the prevalence of *L. tarentolae* was 8.47% in 2021 and 7.69% in 2022, meanwhile the prevalence of *L. infantum* was of 1.7% and 7.69%, respectively (39,40). Regarding *H. turcicus*, only one gecko was positive for *L. infantum* (12). In this study, adult *T. mauritanica* showed a prevalence of 13/49 (26.53%) for *L. tarentolae* and 4/49 (8.16%) for *L. infantum*. When compared to those prior studies (12,39,40), the prevalence of *L. tarentolae* in *T. mauritanica* is higher in Mallorca Island than in the south of Italy respectively (39,40), both endemic regions of canine leishmaniosis. Interestingly, the prevalence of *L. infantum* is of similar magnitude in both areas. More importantly, this is the first study to detect coinfection with *L. tarentolae* and *L. infantum* in adult *T. mauritanica*, showing that such scenarios occur in wild populations.

Such differences of *L. tarentolae* between studies may be accounted for (i) a high density of geckos on the island, which may facilitate parasite transmission; (ii) a high density and distribution of the sandfly vector, which contributes to the spread of *Leishmania*; and (iii) a high prevalence of *Leishmania* infection in dogs, and potentially other mammals inhabiting the island. To the authors knowledge, this is the first study in Spain to examine the prevalence of *L. tarentolae* in reptile species, with findings showing a high prevalence of *L. tarentolae* in adult *T. mauritanica* on Mallorca. The presence of coinfections (3/49) of *L. tarentolae* and *L. infantum* suggests that individual geckos can harbor multiple *Leishmania* species simultaneously, albeit research is still needed in vitro and in vivo regarding its replication and spread.

These findings highlight the potential role of reptiles in the local ecology of *Leishmania* species. While dogs are established reservoirs of *L. infantum* and asymptomatic human and feline infections are common, the high prevalence of *L. tarentolae* and the presence of *L. infantum* in geckos underscores the need to investigate their ecological and epidemiological significance. Further research on additional sensible host species could enhance understanding of *Leishmania* transmission (23), maintenance and alternative hosts (4), contributing to improved strategies for managing human and animal leishmaniosis.

## Conclusion

This study aimed to examine the prevalence of *L. tarentolae* and *L. infantum* in the wild gecko population, as possible reservoir host, on Mallorca Island. The findings contribute to identifying the presence of sympatric *Leishmania* parasites in wildlife that cohabit with dogs and humans in isolated endemic regions. Results show a higher prevalence of *L. tarentolae* when compared to data from mainland Italy (south). Interestingly, this is the first study to detect *L. tarentolae* and *L. infantum* coinfections in *T. mauritanica*, showing that such scenario is possible in wild populations. Further research is required to contextualize these prevalence data alongside those in humans and companion animals. Such research opens the question whether *Leishmania* infection in geckos may contribute to *Leishmania* infection in humans and other mammals or remains confined to reptilian hosts.

## Acknowledgments

The authors wish to acknowledge the technical assistance of Dr. Anna Mercadé in test automation and the group of Prof. Domenico Otranto for their supply of a positive control sample of *L. tarentolae* for the qPCR testing. The authors would like to acknowledge the natural history illustrator Bruna Roqué (www.brunaroque.com) for her contribution with the illustrations of *T. mauritanica* and *H. turcicus*.

## Ethical statement

The capture, euthanasia, and sampling of wild geckos in this study were authorized by the government of the Balearic Island (permit CEP 21/2023).

